# Hydraulic synchrony of spawning sites amongst Earth’s riverine fishes

**DOI:** 10.1101/2021.10.21.464969

**Authors:** Antóin M. O’Sullivan, Alexander M. Morgan, Robert Newbury, Tommi Linnansaari, Barret L. Kurylyk, Jani Helminen, Bernhard Wegscheider, Robert C. Johns, Kurt Samways, Kari I. Alex, R. Allen Curry, Richard A. Cunjak

## Abstract

Earth’s riverine fishes utilize a suite of reproductive guilds, broadly following four guilds: nest guarders, broadcast pelagic spawners, broadcast benthic spawners and nest non-guarders ^1^,^2^, and these guilds utilize different mechanisms to aerate eggs ^3,4^. Globally, river fishes populations are declining^5^, and spawning habitat rehabilitation has become a popular tool to counter these declines^6^. However, there is a lack of understanding as to what classifies suitable spawning habitats for riverine fishes, thereby limiting the efficacy of these efforts and thus the restoration of the target species. Using data from *n* = 220 peer-reviewed papers and examining *n* = 128 unique species, we show the existence of a hydraulic pattern (defined by Froude number (*Fr*), a non-dimensional hydraulic parameter) that characterizes the reproductive guilds of riverine fishes. We found nest guarders, broadcast pelagic spawners, benthic spawners, and nest non-guarders selected sites with mean *Fr* = 0.05, 0.11, 0.22, and 0.28, respectively. Some of the fishes in this study are living fossils, suggesting that that these hydraulic preference patterns may be consistent across time. Our results suggest this hydraulic pattern can guide spawning habitat rehabilitation for all riverine fish species globally in absence of specific spawning habitat information for a species, where resource managers can establish the reproductive guild of the species of interest, and then apply the specific hydraulic requirements (*Fr* range) of that reproductive guild, as presented herein, in the rehabilitation of the target species.

## Main

Throughout the year Earth’s rivers pulse as their fishes spawn to complete their life cycle, thus ensuring the continued success of their respective species. Their selection of spawning sites is based on reproductive guilds, and these broadly (but not exclusively) follow four guilds^1^,^2^: (i) nest builders that guard their nest (nest - guarder), *e*.*g*., smallmouth bass (*Micropterus dolomieu*), (ii) nest builders that do not guard their nest (nest non-guarder), *e*.*g*., Dwarf Ayu (*Plecoglossus altivelis*), (iii) broadcast pelagic spawners – fishes that spawn in the water column (broadcast - pelagic), *e*.*g*., Darling River hardyhead (*Craterocephalus amniculus*) and (iv) broadcast benthic spawners – fishes that broadcast spawn onto substrates (broadcast - benthic)), *e*.*g*., American paddlefish (*Polyodon spathula*). Regardless of the guild, a key to maturation and hatching of these eggs is adequate oxygenation ^3,4^ – see Fig. 1. Nest builders, for instance, fan their nests to aerate their eggs, whereas nest non-guarders will create a nest that modulates river flow to aerate their eggs^7^.

**Fig. 1.**
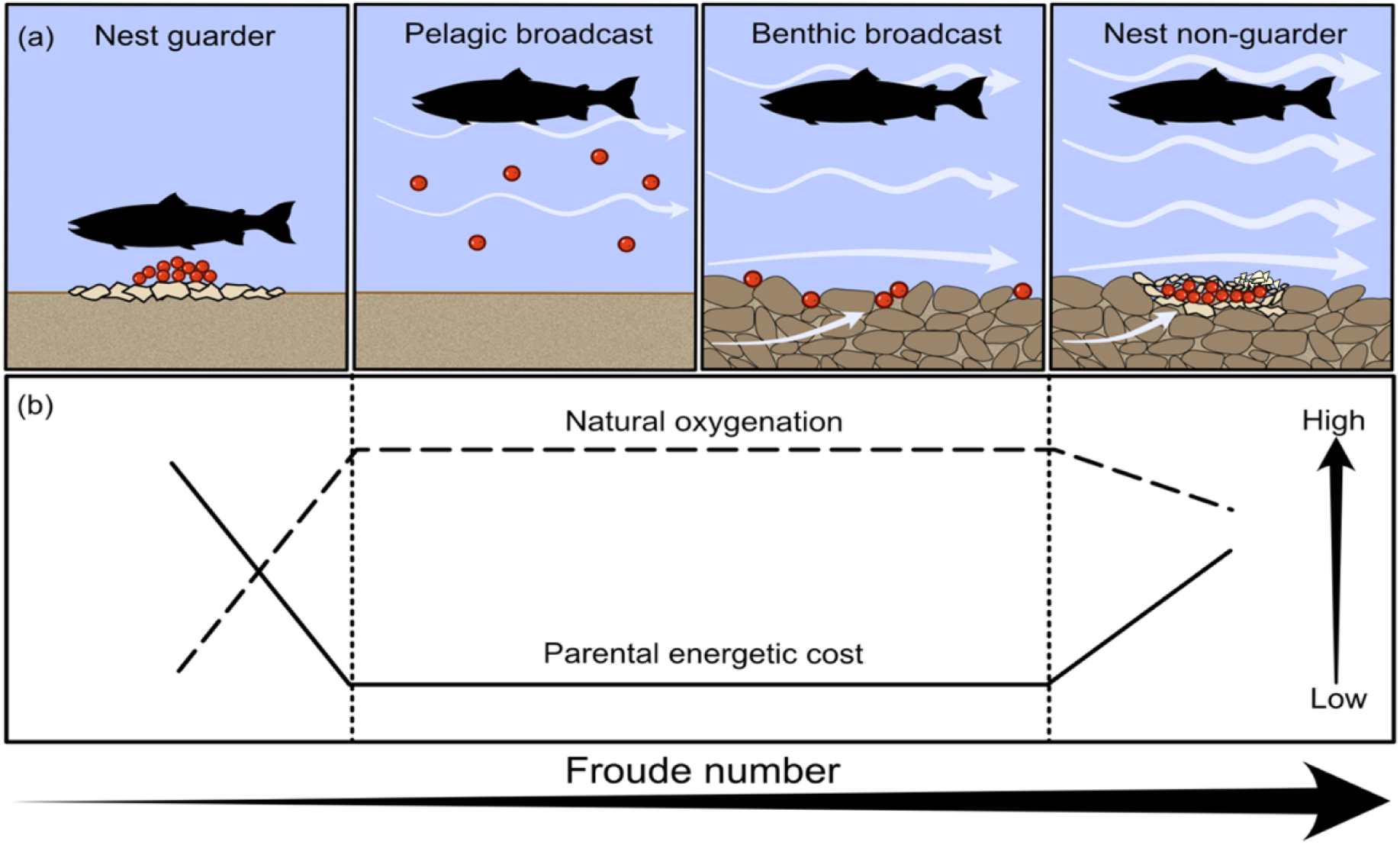
The conceptual model illustrating our hypothesized relationship between global reproductive guilds and Froude number. A schematic of global riverine reproductive guilds (a), and their hypothesized relationship with oxygenation, and parental energetic investment in incubation (b) as a function of Froude number.

Earth’s riverine fishes are experiencing steep declines^5^. The reasons for this are multifaceted and linked to habitat degradation, dams, overharvest, introduction of non-native species, and climate change^8,9^. With these declines, we are on the cusp of losing myriad species and the critical ecosystem functions and services they provide. To counter these declines, governmental and non-profit groups are turning to spawning habitat restoration ^6,10^. While this is a laudable aim, there is a lack of understanding as to what classifies suitable spawning habitats for riverine fishes, and it is infeasible to find this explicitly out for every riverine fish species. This knowledge gap serves to limit the efficacy of these efforts and thus the restoration of the target species. We hypothesize that a hydraulic pattern exists in the spawning habitats of Earth’s riverine fishes, and this pattern can guide spawning habitat rehabilitation in absence of species-specific information. We select Froude number (*Fr*) as a non-dimensional hydraulic parameter that is comparable across all spatial scales, and has shown promise in previous studies examining detailed spawning characteristics (*e*.*g*.,^10,11^). *Fr* is simply a ratio of inertial to gravity forces (see methods), and describes the river’s hydraulic energy regime. We postulate, *a priori*, that low *Fr* values characterize the spawning habitats of nest guarders, where oxygenation of the eggs is provided by the parent. This energetically costly guild motivates the selection of a hydraulic regime that limits energetic expenditure associated with swimming (Fig. 1). Conceptually, as the reproductive guild changes, the hydraulic regime required to aerate the eggs is also predicted to change. For instance, broadcast pelagic spawners that spawn semi-buoyant eggs are hypothesized to select different hydraulic regimes than broadcast benthic spawners that spawn adhesive eggs (Fig. 1). Contrastingly, we hypothesize that nest-non guarders select relatively higher *Fr* values to limit sedimentation accumulation in the nest that would reduce natural oxygenation (Fig. 1). To test our hypotheses, we conducted a meta-analysis of *n* = 220 peer-reviewed papers that describe depth and velocity conditions, and thereby *Fr*, at the spawning sites of *n* = 128 unique riverine fishes across the planet (Fig. 2a and Tables S1 and S2). These data include each of the four reproductive guilds detailed above (Fig. 1 and 2).

**Fig. 2.**
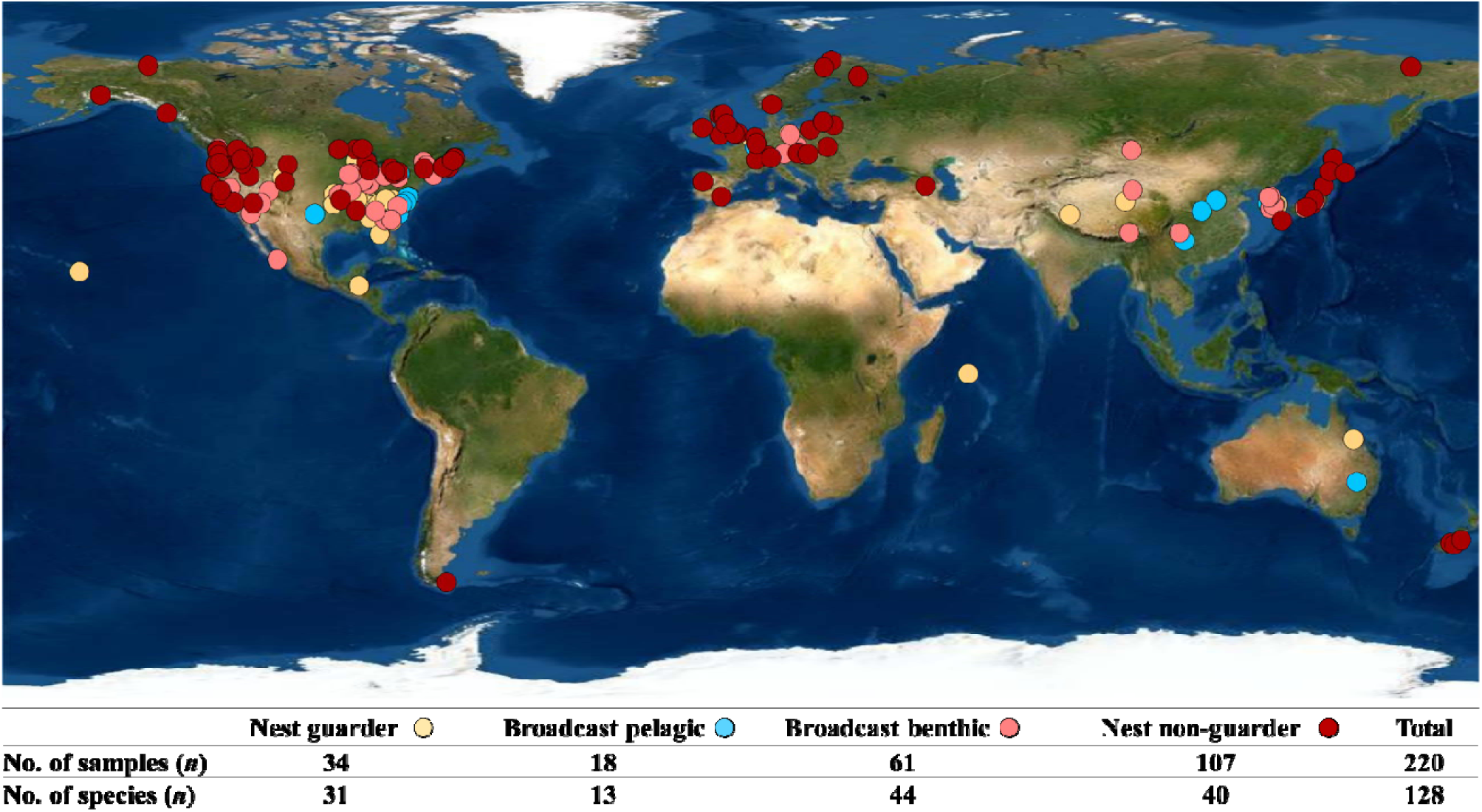
Spatial extent of the data underlying our analysis displaying reproductive guild classification (color), sample size and number of unique species, and the total number of samples for each reproductive guild.

## Results and Discussion

Globally, *Fr* values indicate an association with reproductive guilds that was not evident in depth or velocity (Fig. 3). Nest guarders, broadcast pelagic spawners, benthic spawners, and nest non-guarders selected sites with mean Fr numbers of 0.05, 0.11, 0.22, and 0.28, respectively (Fig. 3). We acknowledge that we observed between-population variability within species in the *Fr* at their spawning location. This means that the *Fr* at selected spawning sites is not necessarily a “crisp” number and some inherent variability is assumed. However, we tested the validity of the observed patterns in *Fr* by examining species specific *Fr* estimates (*i*.*e*., for multiple single species observations, we reduced the parameter of interest to the mean of the total observations), and the results remain the same (see methods and Fig. 3a-c and d-f). As such, our meta-analysis supports the hypothesis, and conceptual model, that a hydraulic pattern, as represented by *Fr*, is apparent from the reproductive guilds of Earth’s riverine fishes (Figs. 1 and 3.).

**Fig. 3.**
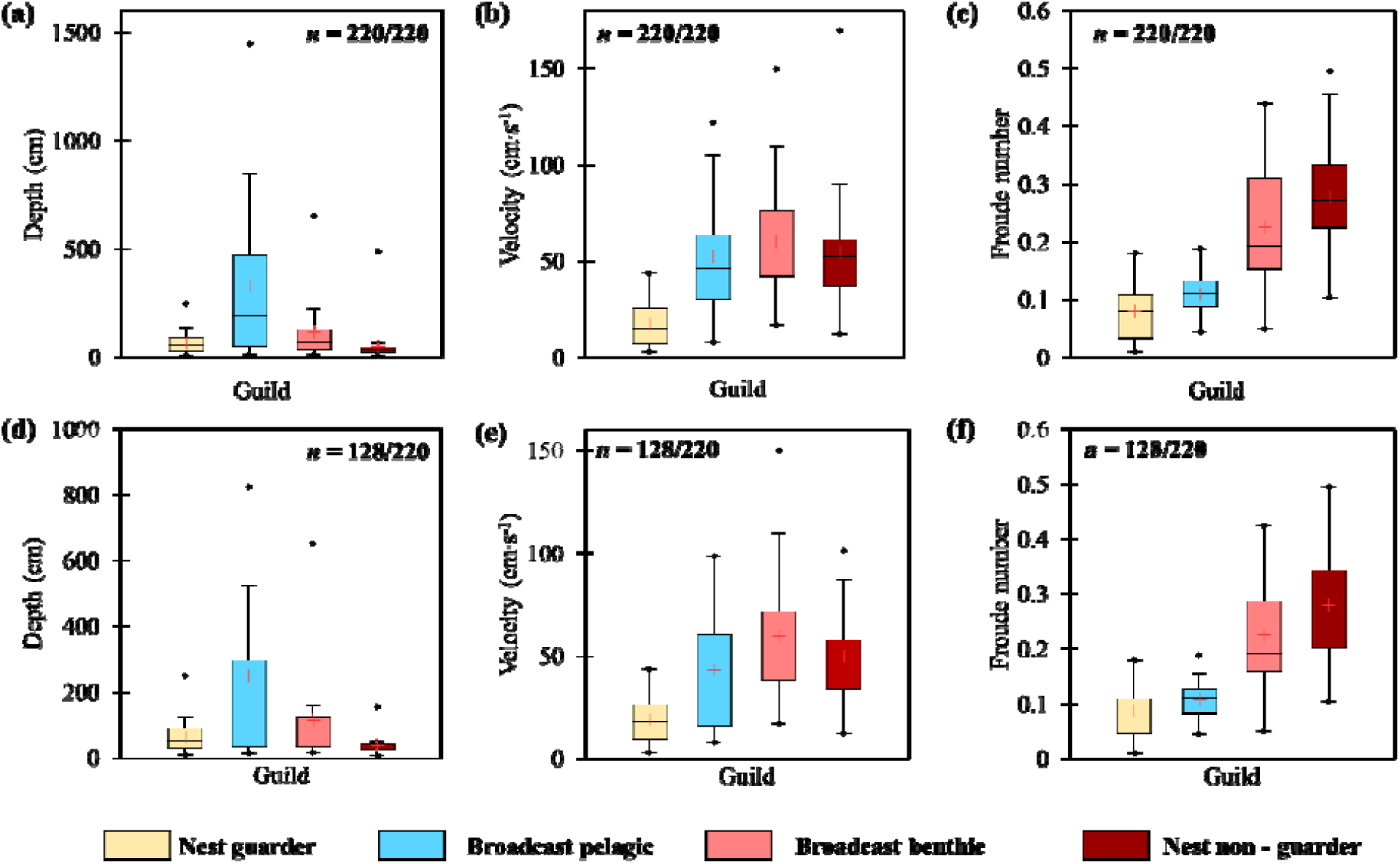
Descriptive statistics results illustrating the relationship between depth, velocity and Froude number values for nest guarders, broadcast pelagic, broadcast benthic, and nest non-guarding reproductive guilds stratified as per our conceptual model (Fig. 1) for the entire data set (*n* = 220) (a-c), and unique species (*n* = 128) (d-f).

Nest guarders selected sites with the lowest *Fr* values observed across the *n* = 128 species examined. Nest guarders, such as the eel tailed catfish (*Tandanus tandanus*), are uniquely characterized in our dataset by a parent that protects and aerates the nest. It is well established that providing parental care is energetically costly, with links to reduced parental growth^12^. Biotic interactions by predators can further exacerbate energetic costs for nest guarders (*e*.*g*., smallmouth bass (*Micropterus dolomieu*) and round goby (*Neogobious melanostomus*) – see ^13^). We investigated the role of *Fr* on energetic expenditure for nest guarders using an integrated Froude number – Strouhal number model (*Fr-St*), where *St* is a dimensionless number that describes oscillating flow mechanisms. Considering nest guarders fan their nest to remove fine sediments and aerate their eggs, we assume *St* = 0.3 as this provides the greatest propulsion to entrain particles and aerate eggs (*e*.*g*., ^14,15^). Examining the relationship between *Fr* and tail-beat frequency (*f*) reveals the energetic expenditure required to obtain optimal *St* – see methods. The *Fr-St* model suggests *Fr* values selected by nest guarders are energetically more efficient than the other reproductive guilds (see *f* values in Table 1). We thus propose the global occurrence of low *Fr* values for nest guarders is linked to energetic conservation of the parent, facilitating removal of fine sediments and aerating eggs in the most energetically efficient manner.

**Table 1.**
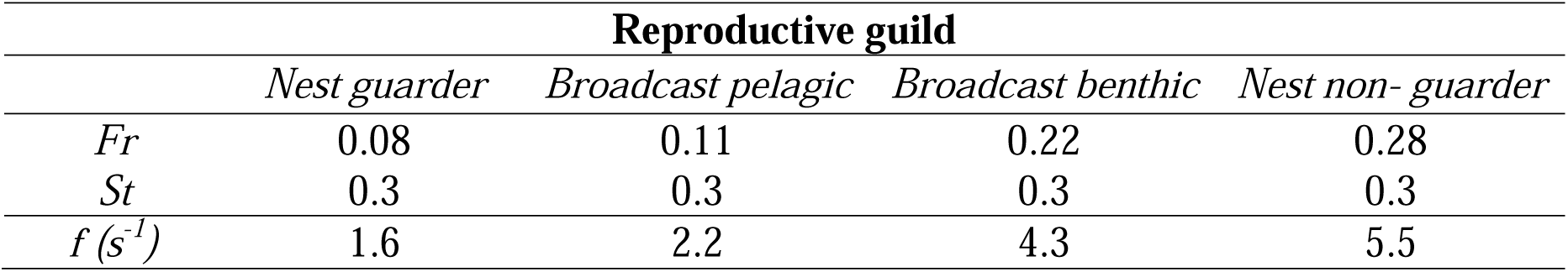
Integrated Froude number (*Fr*) and Strouhal number (*St*) model results for different *Fr* scenarios associated with each reproductive guild quantifying how an increase in *Fr* while retaining an optimal *St* = 0.3^15^, leads to an increase in tail beat frequency *f* (see SI for full model).

The role of hydraulics for the remaining three reproductive guilds is markedly different from nest guarders. When broadcast pelagic spawners spawn in the water column, their eggs can either settle onto vegetation or drift, *e*.*g*., common bream (*Abramis brama*) and black carp (*Mylopharyngodon piceus*), respectively. For species whose eggs are deposited on vegetation, *Fr* values of ∼ 0.11 likely produce an environment that aerates the eggs, while the presence of the vegetation limits shear stress and egg entrainment^16^. For species that spawn directly into the water column, we hypothesize a different link between the egg and the hydraulics. Here, we use striped bass (*Morone saxatilis*) as an example. Upon fertilization, the semi-buoyant striped bass eggs drift in the water column, with eggs becoming denser as they grow^17,18^. In still water, ergo with little drag force, these denser eggs will sink, reducing survival^19^. As such, we suggest the selection of *Fr* ∼ 0.11 for this guild provides adequate flow to entrain the eggs, and as the egg density increases, this hydraulic condition facilitates the egg staying buoyant, *i*.*e*., prevent complete sinking, and allows time for the egg to hatch (Fig. 1).

The hydraulic controls on nest building non-guarders’ site selection have been well studied ^20,21^. Salmonid redds, for instance, are understood to induce hyporheic flow paths (blue arrows Fig. 1) that remove fine sediments and metabolic by-products, while also aerating eggs, and creating a stable thermal environment^22^. These same mechanisms most likely transcend to other nest building non-guarders, such as sea lamprey (*Petromyzon marinus*) – see Table S1. However, less is understood about the role of hydraulics for broadcast benthic spawners, such as the robust redhorse (*Moxostoma robustum*) and alligator gar (*Atractosteus spatula*). Given both nest non-guarders and broadcast benthic spawners select gravel and cobble substrata, and the *Fr* values for both are the highest across the reproductive guilds, we postulate a similar hydraulic mechanism in both instances. Nonetheless, the difference in *Fr* for each guild does suggest subtle mechanistic differences. Broadcast benthic spawners eggs are typically adhesive^1^. The eggs of Asp (*Leuciscus aspius*), for instance, are both negatively buoyant and adhesive^23^. It is also well documented that benthic spawner eggs drop into interstitial voids^6,24^. For broadcast benthic spawners (that spawn on complex bedforms with high hydraulic conductivity (*K*) substrate), we hypothesize the following: (a) *Fr* values ∼ 0.22 induce hyporheic flows in the uppermost section of the substrate^25^; and (b) the adhesive nature of the eggs offsets shear stress and uplift from the bulk flow and hyporheic flow, respectively, thus defining a different guild to exploit the hydraulics in comparison to nest non-guarders (Fig. 1). Similarly, nest non-guarders typically select spawning habitats that are also characterized by complex bedforms with high *K* sediments, and these induce hyporheic flow^26^. However, nest building non-guarders are suggested to select sites with higher *Fr* values (Fig. 3). Nest building non-guarders do not typically have adhesive eggs and will bury their eggs post fertilization in a redd *e*.*g*., brook trout (*Salvelinus fontinalis*) and barbel (*Barbus barbus*). We hypothesize the following for the suggested higher *Fr* values (a) higher *Fr* values will entrain fine sediments during nest construction, and this winnowing is understood to change *K* and thereby down and upwelling flux^22^, and (b) as the eggs of nest building non-guarders are buried, higher *Fr* values are likely required to increase oxygen in the nest^27^.

Some of the fishes in this study can be considered living fossils. The lamprey (*Petromyzontiformes*) and the American paddlefish (*Polyodon spathula*) have existed since the Early Cretaceous^28,29^, as such we hypothesize that that these hydraulic preference patterns with reproductive guilds may be consistent across time. This simple *Fr* model foregoes consideration of many other important factors, including temperature and nutrients^30,31^. For instance, groundwater upwelling may provide both aeration and thermal stability for eggs, thus limiting the requirement to select specific hydraulics, *e*.*g*., ^32^. However, the simple *Fr* model presented herein for four reproductive guilds does provide compelling evidence that the hydraulics that underpin the spawning habitats of the Earth’s fishes are best characterized by *Fr*. The success of utilizing *Fr* values to rehabilitating spawning habitat is already evident in some rivers. Sockeye salmon (*Oncorhynchus nerka*), for instance, have been observed to select spawning habitats with *Fr* = 0.32 (±0.1)^10^. Sockeye spawning habitat in the Okanagan River (British Columbia, Canada) was severely degraded in the 1950’s when > 90 % of the river was straightened and dyked^33^. Between 2014 - 2018, the channelized river was modified with spawning platforms designed to meet 0.2 ≤ *Fr* ≤ 0.4 during the spawning autumn period^34^. These spawning platforms have been utilized by sockeye salmon each year post-construction and have become a critical tool in rehabilitating sockeye salmon. The main finding of the work presented herein is that while it is unfeasible to describe in detail the spawning habitat requirements for all approximately 15 000 freshwater fish species^35^, it may be sufficient to first establish the reproductive guild of the species of interest, and then apply suitable target *Fr* range, as presented herein, in rehabilitation to meet specific hydraulic requirements of the target species. Future conservation and/or restoration projects that target spawning habitat would therefore benefit from utilizing the link between reproductive guild and *Fr* range.

## Methods

### Meta-analysis data search

We conducted searches in multiple article databases using several different search terms to acquire as many peer-reviewed papers as possible. The search terms and database developed for this study are available in the supplementary information as an .xlsx file.

#### Froude number model

Froude number (*Fr*) is a non-dimensional hydrodynamic parameter that is calculated by:

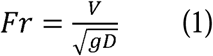

where *V* is water velocity (m·s^-1^), *g* is the gravitational constant = 9.81 m·s^-2^ and *D* is depth (m).

#### Integrated Froude number – Strouhal number model

A Strouhal number (*St*) is a non-dimensional hydrodynamic parameter that describes oscillating flow mechanisms or vortex shedding in a fluid. Myriad experiments have shown that optimal *St* values for swimming animals, defined by maximum propulsive efficiency, range from 0.25 to 0.35 (Taylor *et al*., 2003). For fishes, *St* can be calculated by (Eloy, 2011):

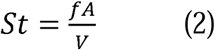

where *f* is the tail-beat frequency (s^-1^), *A* is the peak-to-peak tail amplitude (m), and *V* is water velocity (m·s^-1^).

The *f* term can be isolated by rearranging equations (1) and (2), yielding the following form for the integrated Froude number – Strouhal number mode (*Fr-St*) : Fr.St. -

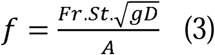

#### Descriptive statistics

We used univariate box plots to examine the relationship each hydraulic parameter and our reproductive guild conceptual model. To account for instances when we had multiple within-species observations, we reduced the observations to one per species to retain equal weights for each species. In such instances, the average of the hydraulic parameters associated with that species were calculated and used for the unique species analysis.

## Acknowledgements

First, the authors would like to thank Lord Pisces. We also thank the Atlantic Salmon Conservation Foundation and New Brunswick Innovation Foundation for funding support. AMM was funded by New Brunswick’s Wildlife Trust Fund. We would also like to thank those scientists whom conducted the studies that underpinned this research.

## Author contributions

Conceptualization: AMO’S; RA Cunjak; Methodology: AMO’S; JH; BW; TL; Investigation: AMO’S; AMM; Visualization: AMO’S; RJ; Funding acquisition: AMO’S, RA Curry, TL; Project administration: AMO’S; Supervision: AMO’S; Writing – original draft: AMO’S; Writing – review & editing: AMO’S; AMM, RN, RJ, TL, BLK, BW, JH, KL, RA Curry, KMS, RN, RA Cunjak

## Materials & Correspondence

Requests can be sent to Antóin M. O’Sullivan.

## Data availability

The supporting data is available as supplementary information.

